# Neural Subspaces Encode Sequential Working Memory, but Neural Sequences Do Not

**DOI:** 10.1101/2025.09.05.674385

**Authors:** Aniol Santo-Angles, Jiarui Yang, Ying Zhou, Wing Kwan Hannah Chu, Grace W Lindsay, Kartik K Sreenivasan

## Abstract

The neural mechanisms of multiple-item working memory are not well understood. In the current study, we address two competing hypotheses about the neural basis of sequential working memory: neural subspaces versus neural sequences. Using broadband MEG data from human participants, we applied dimensionality reduction and multivariate decoding techniques to test whether sequential items are maintained during the retention period through the reactivation of individual items in sequence (neural sequences), or by organizing them into distinct low-dimensional subspaces (neural subspaces). Our results revealed behaviorally relevant, low-dimensional neural subspaces that organized memory representations during the retention period but not during stimulus encoding, supporting the neural subspaces hypothesis. In contrast, we found no evidence of sequential neural replay during the delay period, contrary to predictions from the neural sequences hypothesis. Together, our findings suggest that sequential working memory is maintained through structured geometric organization in low-dimensional representational space, rather than through the sequential reactivation of individual items.

## INTRODUCTION

The neural mechanisms underlying working memory—the ability to hold and manipulate information over short periods—remain incompletely understood, particularly how the brain maintains multiple items simultaneously without interference (D’Esposito 2007; D’Esposito and Postle 2015). Classical theories based on single-neuron recordings propose that working memory relies on persistent firing, with multiple items maintained by distinct neuron populations, each encoding a specific item (Goldman-Rakic 1995; Riley and Constantinidis 2015). However, this view has been challenged by findings that the same neurons often exhibit transient bursts of activity rather than sustained firing. One such proposal is that multiple items are maintained by temporally segregated gamma bursts representing different items within overlapping neural populations, with alpha/beta oscillations regulating selective reactivation and suppression of memory representations (Lundqvist et al. 2016, 2018; Liljefors et al. 2023). Another line of work suggests that working memory relies on the interplay between activity-dependent and activity-silent mechanisms, such as short-term synaptic plasticity, enabling the storage of multiple items in latent states that can be selectively reactivated when needed—thus minimizing mutual interference (Rose et al. 2016; Barbosa et al. 2020; Stokes 2015; Panichello et al. 2024). An emerging view focused on population-level analyses of multiple neuron recordings have reframed working memory as an emergent property of dynamic activity patterns across neural ensembles, where an heterogeneous neural dynamics at the single neurons coexist with the stable representations of memory contents in a low dimensional space (Cueva et al. 2020; Murray et al. 2017; Spaak et al. 2017). Recent work has focused on characterizing the representational geometry of this low-dimensional subspace. In multi-item working memory tasks, individual items are maintained during the retention period in distinct subspaces, which helps minimize interference and preserve behaviorally relevant information (Panichello and Buschman 2021; Tian et al. 2024; Xie et al. 2022). These low-dimensional subspaces not only capture stable memory representations but also reflect ongoing cognitive operations. When two items are held in memory simultaneously, their representations occupy quasi-orthogonal subspaces. Once one item is prioritized and the other becomes irrelevant, the subspaces reorganize, becoming more aligned or parallel (Panichello and Buschman 2021). Similarly, in sequential working memory tasks, the identity of each item is encoded in a distinct subspace according to its ordinal position in the sequence (Xie et al. 2022). If subjects are cued to recall the sequence in reverse order, memory representations swap between subspaces during the retention period to reflect the updated sequence (Tian et al. 2024). While these studies focused on electrophysiological recordings in non-human primates, human neuroimaging research has similarly shown that large-scale brain dynamics evolve within a low-dimensional subspace (Shine, Breakspear, et al. 2019; Shine, Hearne, et al. 2019; Rué-Queralt et al. 2021; Gao, Mishne, and Scheinost 2021). Representational geometry has been extensively studied through representational similarity analysis (Kriegeskorte, Mur, and Bandettini 2008; Kriegeskorte and Kievit 2013; Kriegeskorte and Diedrichsen 2019), which demonstrates that the distances between high-dimensional multivariate neural activity patterns elicited by different sensory inputs reflect information processing (Diedrichsen and Kriegeskorte 2017). Yet, studying neural geometry by leveraging low-dimensional subspaces remains a promising and relatively underexplored direction in human neuroimaging, particularly in the domain of working memory (Kriegeskorte and Wei 2021). In an fMRI study with a single-item working memory task, (Li and Curtis 2023) found that low-dimensional subspaces encoded spatial stimuli in a topologically organized manner within visual and parietal cortices, while this organization was absent in prefrontal areas. In another fMRI study, (Xu 2024) reported that posterior cortical areas orthogonalize memory representations of targets and distractors to avoid interference. These studies suggest that the human brain may leverage low-dimensional neural geometry to separate and maintain memory representations. Nevertheless, it remains unclear whether multiple working memory contents—whether presented simultaneously or sequentially—are also represented in orthogonal subspaces, whether this neural geometry is behaviorally relevant, and whether such neural coding can be detected using non-invasive, large-scale brain imaging techniques like MEG.

Another possibility is that multiple memory contents are embedded within the dynamic patterns of neural activity. This notion is particularly relevant in the context of sequential working memory, where the ordinal position of each item may be encoded through the temporal structure of transient neural reactivations. One compelling example of such temporally structured activity is hippocampal replay, an offline process occurring during sleep or restful wakefulness, in which previously experienced sequences are spontaneously reactivated in either forward or reverse order (Diba and Buzsáki 2007; Foster and Wilson 2006; Gupta et al. 2010). Hippocampal replay has also been implicated in sequence learning, where it is thought to support the extraction and consolidation of temporal structure across events (Davachi and DuBrow 2015; Tacikowski et al. 2024). Although neural replay has usually been studied with intracranial neuronal recordings, recent studies demonstrated that neural replay could also be studied in MEG data with the combination of decoding analysis and linear modelling (Liu, Dolan, et al. 2021). These studies revealed human neural replay of task-relevant sequences deploying at compressed timescales during resting-state periods (Kurth-Nelson et al. 2016; Liu et al. 2019; Liu, Mattar, et al. 2021). However, this approach has not yet been extended to examine online replay, defined as the rapid reactivation of memory sequences during working memory maintenance. In this context, the theta–gamma coding hypothesis of sequential working memory proposes that temporal organization relies on oscillatory phase coding mechanisms, particularly through interactions between theta and gamma rhythms (Jensen and Lisman 2005; Lisman and Idiart 1995; Lisman and Jensen 2013). According to this model, each item in a sequence is represented at a distinct phase of a theta cycle, with item-specific gamma bursts occurring at consistent theta phases, allowing the entire sequence to be maintained within a single theta cycle. Empirical support for this mechanism comes from M/EEG studies showing that gamma activity is systematically modulated by theta phase during the encoding and maintenance of sequential items in working memory, particularly in the hippocampus and prefrontal areas (Bahramisharif et al. 2018; Axmacher et al. 2010; Heusser et al. 2016; Hsieh, Ekstrom, and Ranganath 2011; Borderie et al. 2024). Moreover, the temporal phase patterns observed during memory retrieval and memory encoding of sequences are similar, suggesting that temporal information is preserved during the retention period (Michelmann, Bowman, and Hanslmayr 2016). However, the evidence for online neural replay during the maintenance period of sequential working memory is mixed. An EEG study using a temporal response function (TRF) approach reported rapid backward reactivation of memory items during the delay (Huang et al. 2018). In contrast, an MEG decoding study found no evidence of sequential replay; instead, neural activity was dominated by a single prioritized item, typically the one that had been more weakly encoded (Jafarpour et al. 2017). These findings suggest that while replay may support memory maintenance, its occurrence during the delay period remains uncertain. Hence, evidence of online neural replay during the retention interval of sequential working memory tasks remains scarce.

In the current study, we tested two complementary (but not mutually exclusive) hypotheses regarding the neural representation of sequential working memory (Figure 1A). The neural subspaces hypothesis proposes that memory items are encoded and maintained in distinct, orthogonal neural subspaces, each representing a specific ordinal position. In contrast, the neural sequences hypothesis suggests that sequential information is encoded through the temporal dynamics of neural reactivations, reflecting online replay of memory items. Testing both hypotheses in the same dataset enables a direct, controlled comparison of competing neural coding models for sequential working memory, reducing confounds and revealing potential interactions.

**Figure 1.**
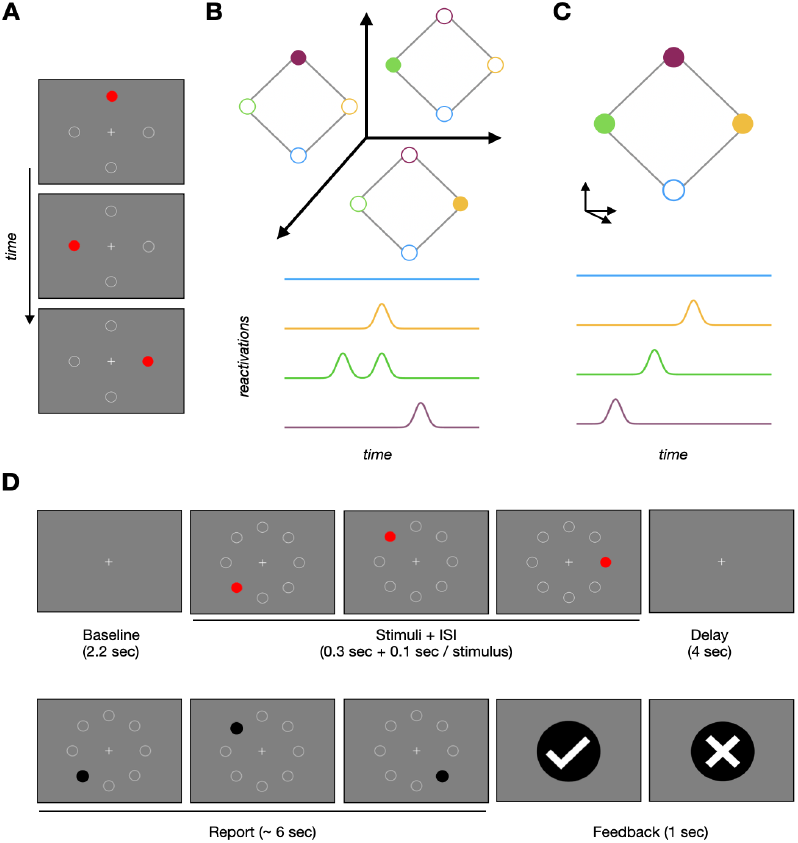
Conceptualization of the study hypotheses about the neural code of sequential working memory. A) Example of a stimulus sequence where three red dots appear in succession (top to bottom) at one of four possible spatial locations (indicated by a white circle as a placeholder). B) Neural subspaces hypothesis: Memory representations are encoded in rank-specific ring attractor models. Different stimuli are encoded simultaneously without interference since rank-specific rings lie in orthogonal subspaces. C) Neural sequence hypothesis: Memory representations are encoded in a ring attractor model embedded in a n-dimensional space. Rank information, i.e., the ordinal position within the sequence, is encoded in the dynamics, where the timing of reactivations during delay period follow the order of stimuli presentation, and the time lag between reactivations remain constant. D) Design of the sequential WM task.

## RESULTS

### Task description and behavioral performance

Seventeen healthy participants performed a sequential working memory task (Figure 1B) while their brain activity was recorded using magnetoencephalography (MEG). During the task, participants observed a sequence of stimuli and were required to reproduce the sequence accurately. Participants performed well, successfully reproducing the presented sequence on 83 ± 19% of trials across sessions, significantly above chance according to a binomial test (p < 1e-15 for all subjects). A linear regression analysis showed no significant effects of sequence length, session, or their interaction on task performance (all p > 0.05), suggesting consistent performance across different experimental conditions and sessions. This consistency justified pooling trials across sessions for subsequent analyses. After preprocessing MEG data, an average of 267 ± 41 trials per subject were included in the analysis, with 221 ± 50 trials being correct responses, comprising 126 ± 15 correct responses for sequences of length-3 and 96 ± 14 correct responses for sequences of length-4.

### Geometric analysis of neural subspaces

To test the hypothesis that sequential working memory contents were encoded in neural subspaces, we adapted the analysis pipeline described in previous work (Panichello and Buschman 2021; Xie et al. 2022) for application to MEG data. Because signals from a common neural source can be detected by multiple distant sensors (Hari et al. 2017) potentially introducing spurious correlations that artifactually align neural subspaces in sensor space, we performed all analyses in source space. Specifically, we examined the geometry of neural subspaces using source-reconstructed data from a subset of 15 participants for whom individual MRI scans were available. The source data were parcellated into 200 cortical regions (Schaefer et al. 2018), and corrected for signal leakage using symmetric orthogonalization (Colclough et al. 2015). Neural subspaces for each ordinal position in the sequence were defined by projecting subject-specific condition-by-channel matrices of neural activity—where each condition represented a specific stimulus location at a given ordinal position in the sequence—onto their leading three principal components (Figure S1 in Supplementary Material). The degree of alignment between subspaces was quantified using two complementary metrics: the principal angle (PA) and the variance accounted for (VAF) between subspaces (see Methods for details). During the delay period, subspaces corresponding to different ordinal positions were significantly more orthogonal —that is, less overlapping or more discriminable— than expected by chance. This effect was observed for both sequence lengths (3 and 4 items) and held relative to two baselines: a within-subspace comparison and a surrogate-based null distribution. The pattern was consistent across different alignment measures (Figure 2A). In contrast, no consistent subspace orthogonality emerged during the stimulus presentation period. The behavioral relevance of neural subspaces was assessed by comparing trials with correct and incorrect responses after controlling for the imbalanced number of trials (see Methods for details). Incorrect trials showed significantly less orthogonal (i.e., more overlapping) neural subspaces during the delay period (Figure 2B). No significant differences were observed during the stimulus period. For length 4 trials, these differences persisted throughout the entire delay. In contrast, for length 3 trials, the differences were more transient, appearing briefly after stimulus offset and not maintained across the full delay. To further characterize the geometry of neural subspaces, we quantified the separability of memory representations within the subspace of each ordinal position using two metrics: the euclidean distance between stimulus representations, and the volume spanned by those representations in the subspace. Overall, we found no significant effect of performance on separability, except for volume in 4-item trials, where subspace volume was significantly greater for correct compared to incorrect trials (Figure S2). During the delay period of trials of length 4, but not length 3, the volume of stimulus representations in the neural subspaces was significantly greater in correct than incorrect trials. No differences were observed when separability was measured with the euclidean distance. To assess the effect of memory load, we compared geometric variables between trials of length 3 and length 4 (Figure S3). Subspaces were generally more orthogonal (less aligned) for length-3 trials, though this difference was not consistently significant and only reached significance in disconnected time windows.

**Figure 2.**
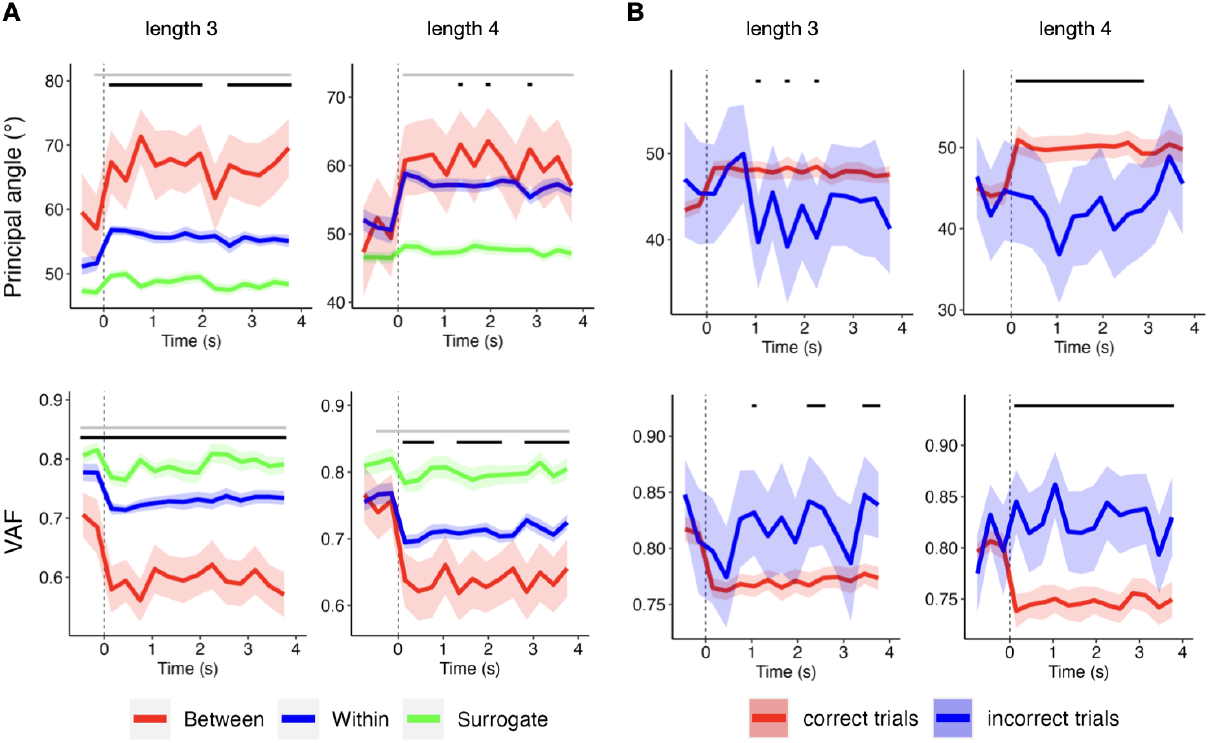
Geometric analysis of neural subspaces. Alignment between rank-specific subspaces, quantified using the principal angle (PA, top row) and variance accounted for (VAF, bottom row). A) Comparison of alignment measures for correct trials: empirical between-subspaces (red), surrogate between-subspaces (green), and within-subspaces (blue). B) Comparison of empirical between-subspace alignment in correct (red) versus incorrect trials (blue). X-axis shows time locked at delay onset (vertical dashed line). Black (grey) horizontal lines on top indicate time segments with FDR-corrected significant differences between the variables represented by red vs blue (red vs green) colors. Shaded areas indicate the standard error of the mean across subjects.

The dimensionality of neural subspaces was evaluated using the eigenvalue spectrum of the data covariance (Figure S4). In trials of length 3 (4), the leading three principal components explained 61 ± 5% (66 ± 5%) of the variance during delay period, and 67±8% (72±8%) during stimuli period. In contrast, when PCA was applied to surrogate data—constructed by independently shuffling rows within each channel—the variance explained by the leading 3 (5) components dropped to 31±0.6% (27±0.4%) for both periods, indicating that the low-dimensional structure of neural subspaces is not preserved in the shuffled controls. The contributions of different brain networks to the neural subspaces were evaluated by projecting the leading three eigenvectors onto brain maps (Figure 3). These maps were then averaged across time windows and subjects to identify consistent spatial patterns of involvement. To quantify network-specific contributions, eigenvector values were averaged across parcels within each network. Results showed that the stimulus and delay periods exhibited a similar distribution of parcel and network engagement. Among the networks, the visual network contributed the most, accounting for 31% of the total magnitude across all parcels, followed by the dorsal attentional network (20%), default mode network (16%), limbic network (13%), somatomotor network (10%), control network (7%), and salience/ventral attentional network (3%).

**Figure 3.**
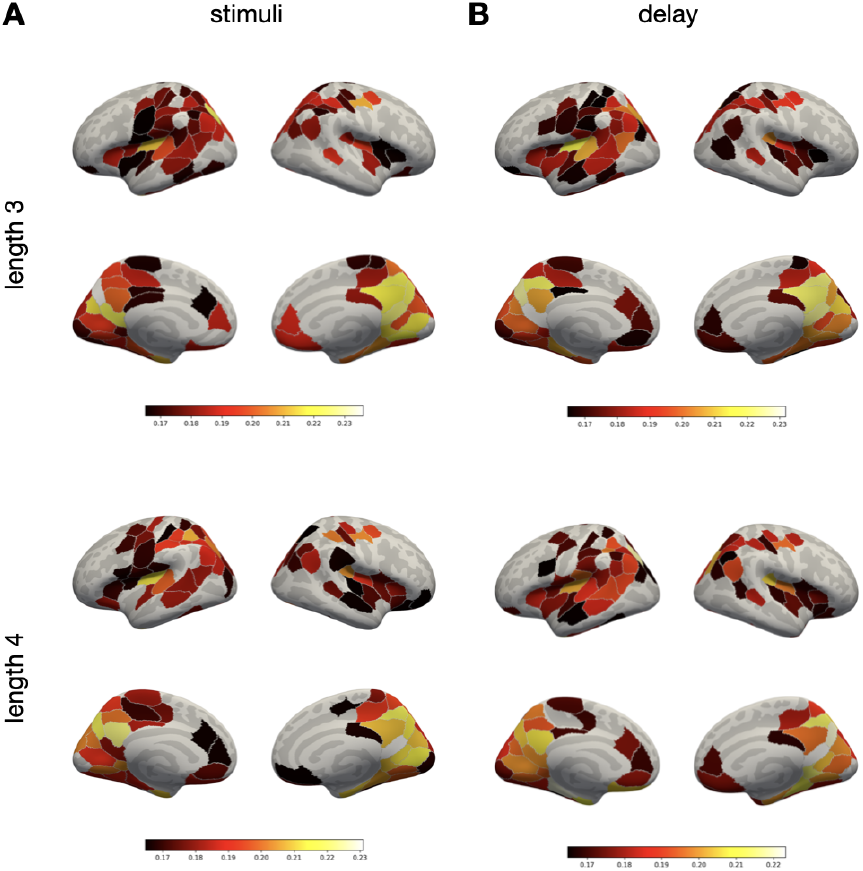
Brain maps of the eigenvectors defining the neural subspaces, averaged across time windows and subjects, during A) stimuli period and B) delay period, for trials of length 3 (top row) and length 4 (bottom). For visualization, maps are thresholded at the top 50% of values to highlight regions with the strongest contributions.

Supplementary analyses revealed no overall qualitative changes in the main findings when neural subspaces were defined using the leading six or eight principal components, confirming the robustness and reliability of our results across different subspace dimensionalities (Figures S5 and S6, Supplementary Material). However, when eight principal components were used, the alignment between subspaces was no longer significantly more orthogonal than within-subspaces, although it remained significantly greater than that observed in surrogate data. This likely reflects the reduced representativeness of the two-dimensional planes used to compute principal angles (PA) and variance accounted for (VAF) in higher-dimensional spaces (see Methods for details). Additionally, we found that the effect of performance on subspace separability—measured through distance and volume—became significant across all trial lengths when eight principal components were used (Figures S6, Supplementary Material). We also observed that subspace volume systematically decreased with increasing numbers of principal components. This suggests that increasing the dimensionality of the subspace does not lead to a greater spread of the data but instead incorporates additional dimensions that contribute relatively little variance. As a result, the overall volume becomes compressed, reflecting the diminishing returns of adding higher-order components that capture less structure in the data. The second supplementary analysis, when the data matrix of neural activity was defined by grouping spatial locations to increase the signal-to-noise ratio, revealed no qualitative changes in the main results, although the effect of performance on the volume of the neural subspace for trials of length 4 was only significant within a single time window, rather than sustained throughout the delay period (Figure S7). Finally, when the geometric analysis was performed in sensor space, neural subspaces were significantly more orthogonal than expected by chance. However, the effect of behavioral performance observed in source space did not persist, suggesting that sensor space may obscure behaviorally relevant neural geometry due to signal mixing (Figure S8).

### Sequence effects of memory representations

To test the hypothesis that sequential information was encoded in the dynamics of memory reactivations, we applied the temporally delayed linear modelling (TDLM) (Liu, Dolan, et al. 2021). We trained linear classifiers on sensor MEG data (n=17) and source-reconstructed data (n=15) to identify spatial locations during stimulus presentation in both the localizer and working memory tasks. Classifiers achieved significant cross-validated classification accuracy spanning through most of stimulus period, peaking around 150 ms after stimulus onset during localizer (sensor: peak = 160 ± 125, AUC = 0.56 ± 0.02, p < 1e-5; source: peak = 140 ± 50 ms, AUC = 0.63 ± 0.05, p < 1e-5) and WM task (sensor: peak=150 ± 90 ms, AUC = 0.66 ± 0.05, p < 1e-5; source: peak=155 ± 85, AUC = 0.59 ± 0.03, p < 1e-5) (Figure 4 and S4). Since sensor-level classifiers outperformed source-level classifiers, the analysis of sequence effects was performed on sensor data. The most accurate classifiers were then used to construct a decoded state space of memory representations by applying the classifier weights to delay period data (see Methods section for the group-level selection of the most accurate classifier). This process generated a matrix representing decoded state space (time by spatial locations), encoding the probability of memory reactivations throughout the delay period. We also computed the decoded state space of sensory representations by repeating the same process on WM task stimuli period data. To quantify sequence effects at the subject level, two complementary approaches were employed. The first approach involved calculating sequence effects at the trial level and averaging these effects across all trials (Figure 4B) (Schwartenbeck et al. 2023). The second approach concatenated the decoded state space across trials, allowing sequence effects to be computed collectively for all trials (Figure 4C). This method aligns with the longer timescales for which TDLM was originally developed, specifically to investigate neural replay during resting state (Liu et al. 2019; Kurth-Nelson et al. 2016), and mitigates finite sample effects inherent to trial-level analyses. Both approaches yielded consistent results. During the stimulus presentation, TDLM identified a significant forward sequence effect, peaking at a time lag of 400 ms. This timing matched the expected dynamics given the 300 ms stimulus presentation and 100 ms interstimulus interval (ISI) (Figure 4B and 5C first and second rows). These results were consistent across decoded state spaces computed using classifiers trained on either WM stimuli period data (stim-to-stim) or localizer data (localizer-to-stim). During the delay period, we observed no evidence of forward or backward sequence effects, regardless of whether classifiers were trained on WM stimuli period data (stim-to-delay) or localizer data (localizer-to-delay) (Figure 4B and 5C, third and fourth rows). Since classifier performance was reduced for discriminating between nearby locations (Figure S10), we repeated the analysis of sequence effects, excluding trials in which the sequence contained locations separated by only one step (e.g., the sequence [1, 2, 4] was excluded, but not [1, 3, 5]). This analysis confirmed our previous findings (Figure S11). In summary, we found no evidence of neural sequences of memory reactivations during delay period, either at trial- or subject-level.

**Figure 4.**
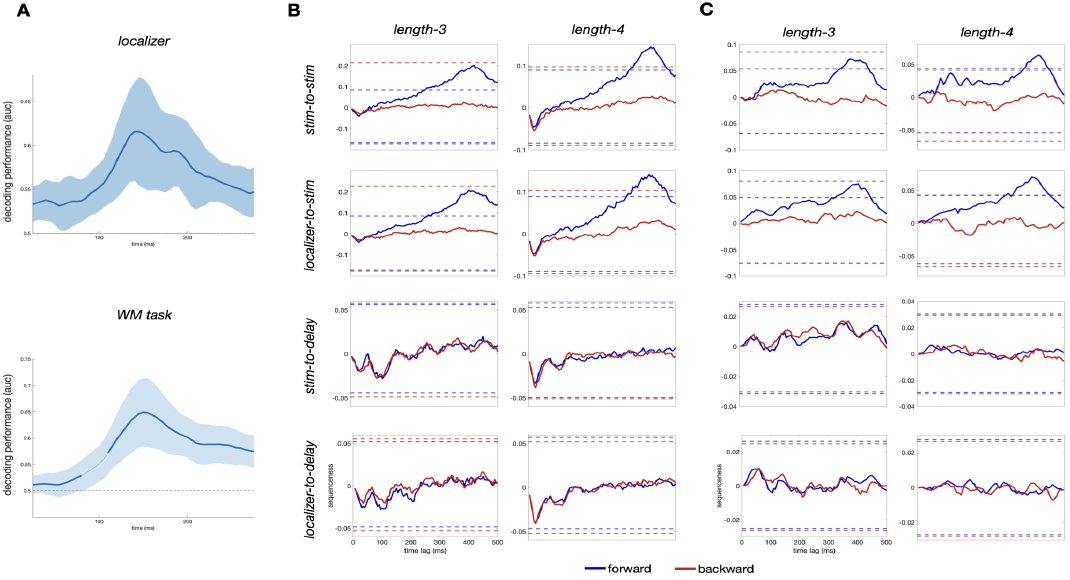
(A) Decoding performance for stimulus location during stimulus presentation in the localizer task (top) and sequential WM task (bottom), time-locked to stimulus onset. The solid lines and shaded areas represent the mean and standard deviation of the area under the curve (AUC) across subjects. Thicker lines indicate decoding performance significantly above chance (dashed line at AUC = 0.5). (B-C) Sequenceness effect at (B) trial-level and (C) subject-level. Rows (top to bottom) show: a) stim-to-stim: classifier trained on WM-stimuli data and applied on WM-stimuli data to compute decoded state space, b) localizer-to-stim: classifier trained on localizer data and applied on WM-stimuli data, c) stim-to-delay: classifier trained on WM-stimuli data and applied on WM-delay data, and d) localizer-to-delay: classifier trained on localizer data and applied on WM-delay data. Solid lines represent forward (blue) or backward (red) sequenceness effects averaged across subjects, dashed lines indicate the upper and lower thresholds for statistical significance.

## DISCUSSION

In this study, we tested two competing hypotheses about the neural mechanisms underlying sequential working memory: neural subspaces versus neural sequences. Our results support the neural subspace hypothesis, showing that memory items are maintained in low-dimensional, behaviorally relevant neural subspaces. In contrast, we found no evidence supporting the sequential reactivation of memory contents.

Our findings on neural subspaces replicate previous results from neuronal recordings in non-human primates, demonstrating quasi-orthogonal subspaces during the retention period. We observed principal angle values of approximately 70 degrees for trials of length 3 and 60 degrees for trials of length 4—comparable to the 70–80 degree range reported in non-human primates (Xie et al. 2022; Panichello and Buschman 2021; Tian et al. 2024). Notably, the patterns of subspace alignment differed between the stimuli and retention periods. Specifically, during the retention period, subspaces were significantly more orthogonal (i.e., less aligned) than expected by chance. This effect was robust across different methods of estimating chance levels, trial lengths (though more sustained in length-4 trials), and alignment metrics. In contrast, during the stimuli period, subspaces were generally not different from chance—except for trials of length 3, where increased alignment was observed using the VAF metric. Interestingly, the brain maps reflecting cortical parcel contributions to the neural subspaces were remarkably similar across stimuli and retention periods (Figure 3). These maps revealed a consistent spatial pattern involving occipital and parietal cortices—recruiting visual and dorsal attention networks— as well as left-lateralized prefrontal and temporal regions, brain areas widely associated with working memory (Sreenivasan and D’Esposito 2019). Taken together, these findings suggest that encoding and maintenance engage overlapping brain networks, in line with the sensory recruitment hypothesis (Pasternak and Greenlee 2005; D’Esposito 2007; Sreenivasan et al. 2014). However, the computations supported by these regions appear to differ qualitatively across phases: neural subspaces emerge only during the maintenance period, not during stimulus encoding. Consistent with this idea, (Kwak and Curtis 2022) demonstrated that working memory representations of orientation and motion direction in visual and parietal cortices are encoded in an abstract, line-like format that preserves only task-relevant information. Furthermore, prior work has shown that sensory representations are transformed into memory representations via orthogonalization through rotational dynamics, allowing perceptual and mnemonic codes to be segregated within overlapping neural populations (Libby and Buschman 2021).

The geometry of subspaces in our data was behaviorally relevant, consistent with previous studies. (Xie et al. 2022) showed that subspace geometry predicts systematic memory errors: greater overlap between adjacent rank subspaces led to transposition errors, while ring-like structure within subspaces accounted for confusions between nearby spatial locations. (Panichello and Buschman 2021) also observed more overlapping subspaces in error trials. We replicated this pattern, observing increased overlap between subspaces in incorrect trials. This suggests that orthogonal—or at least sufficiently distinct—neural subspaces are necessary to keep memory representations discriminable. In contrast, excessive alignment between subspaces reduces this discriminability, making representations more confusable and thus more prone to behavioral errors. These results support the idea that distortions in neural subspace geometry may underlie failures in working memory performance. Regarding the separability of stimulus representations, when subspaces were computed using 3 PCs, we initially found no differences between correct and incorrect trials, except for the volume metric in trials of length 4, where correct trials exhibited greater volume than incorrect trials. The selective effect on volume may suggest that subspaces under higher memory load exhibit greater nonlinearity in their geometry. Unlike pairwise distances, volume reflects the overall shape and spread of the representation, capturing more complex spatial configurations. For example, a flat and a curved (e.g., bent) ring structure with equally spaced item representations may yield similar euclidean distances between points, but the curved structure would enclose a larger volume when measured using the convex hull. The fact that this effect emerged only in trials of length 4 may reflect a memory load–dependent distortion of the representational geometry. Consistent with this interpretation, principal angles were smaller in length-4 than length-3 correct trials. Extending the analysis to subspaces computed with 8 PCs revealed additional effects: volume differences also appeared in length-3 trials, and pairwise distance differences emerged in length-4 trials. These findings indicate that higher-dimensional subspaces reveal aspects of the representational geometry that are less apparent in lower-dimensional projections. This could reflect nonlinear distortions, but it could also indicate that memory representations are fundamentally linear yet occupy a higher-dimensional space that is missed when projecting onto only 3 PCs. Together, these findings suggest that memory load may influence both the dimensionality of subspaces and the degree of nonlinearity in their geometry, consistent with prior work linking greater task complexity to more nonlinear representational structures (Fortunato et al. 2024; Shine, Hearne, et al. 2019), although the proposed link between memory load and nonlinear distortions in subspace geometry remains speculative and should be tested experimentally.

Contrary to predictions from the neural sequences hypothesis, we did not observe evidence of sequential reactivation during the retention period. We applied a validated method, Temporally Delayed Linear Modelling (TDLM) (Liu, Dolan, et al. 2021), to estimate neural sequences in broadband MEG data. This approach has reliably detected neural sequence replay during resting-state periods (Kurth-Nelson et al. 2016; Liu et al. 2019) as well as task-related sequential reactivation (Schwartenbeck et al. 2023), demonstrating its sensitivity and robustness in identifying temporally structured neural activity. Our null finding was consistent across multiple analysis strategies. We trained classifiers both on a localizer task, which contained no inherent sequence structure, and on stimulus presentations during the working memory task, achieving significant decoding performance in both cases. The decoded state space successfully identified expected sequences of reactivation during stimulus presentation, confirming the validity of our classifiers—even when trained on the localizer data and tested on WM task data. To further validate our results, we adapted the TDLM pipeline to control for finite sample effects at the trial level and concatenated retention periods across trials to increase statistical power. Despite these efforts, no evidence of sequential reactivation emerged during the retention period. Our results align with those of (Jafarpour et al. 2017), who also found no evidence of neural replay during the retention period in an MEG study of sequential working memory. Instead, their decoding analysis revealed that neural activity during the delay was dominated by a single item, regardless of its original position in the sequence or category. This prioritized item tended to be one that had been more weakly encoded during the stimulus period, suggesting that attention may selectively enhance the maintenance of less robust memory traces, rather than supporting uniform sequential replay. In contrast, our findings diverge from those of (Huang et al. 2018), who reported evidence of neural replay during the retention period in a sequential working memory task. Using a temporal response function (TRF) approach, they tracked item-specific reactivations by presenting task-irrelevant visual probes—each uniquely colored and previously associated with a memory item—and modeling alpha-band responses to these flickering stimuli. Their results revealed fast, reverse-order replay of items within brief (200–400 ms) time windows. While this approach offers valuable insights, its reliance on rhythmic luminance introduces interpretive challenges, as the observed reactivations may reflect stimulus-evoked responses rather than spontaneous memory processes. Indeed, prior work has shown that flicker-based “pinging” can trigger retrieval of latent memory representations (Barbosa, Lozano-Soldevilla, and Compte 2021; Wolff et al. 2017), reorganizing working memory contents by decreasing neural dynamics (Yang, He, and Cai 2025), raising the possibility that the reported sequential reactivations were externally driven. In this context, decoding-based approaches, which track spontaneous reactivations of memory content during the delay period, may offer a more direct test of whether sequential replay underlies working memory maintenance (Liu et al. 2022; Liu, Dolan, et al. 2021). Our null findings regarding neural sequences also resonate with behavioral and neuroimaging studies suggesting that working memory compresses sequential information using abstract geometric and numerical primitives (Amalric et al. 2017; Wang et al. 2019; Al Roumi et al. 2021). According to this view, rather than storing and replaying individual items in sequence, the brain encodes higher-order structure—such as symmetry, rotation, or numerical rules—that generates the entire sequence. This form of mental compression could reduce the need for item-by-item reactivation during the retention period, potentially explaining the absence of detectable neural replay in our data. However, if rule-based compression were the only mechanism supporting sequential working memory, the neural subspace patterns we observed would likely not emerge.

Our null findings are somewhat unexpected in light of the theta–gamma coding model of sequential working memory, which proposes that individual memory items are represented by gamma-band activity nested within distinct phases of the theta cycle (Lisman and Jensen 2013). This model is supported by evidence of cross-frequency coupling modulation during both the encoding and maintenance periods (Bahramisharif et al. 2018; Axmacher et al. 2010; Heusser et al. 2016; Hsieh, Ekstrom, and Ranganath 2011; Borderie et al. 2024), showing that the temporal structure of theta–gamma interactions plays a critical role in organizing sequential information in working memory (Roux and Uhlhaas 2014; Davachi and DuBrow 2015). However, a recent study challenged the main prediction of sequential reactivation of memory representations during the maintenance period. (Liebe et al. 2025) recorded spikes and LFPs in the human medial temporal lobe and found increased theta power and spike-phase locking for remembered items. While firing phase reflected item position in correct trials, the sequence of firing phases did not match item order, consistent with our null finding on neural sequences. Taken together with previous work, an alternative interpretation is that theta–gamma cross-frequency coupling observed during sequential working memory tasks, traditionally interpreted as evidence of sequential reactivation of memory representations phase-locked to theta oscillations, may instead reflect that ordinal position information is encoded in subspaces via the phase of low-frequency activity, without necessitating sequential reactivations. Previous work, including our own, has shown that multi-item working memory contents are maintained in quasi-orthogonal subspaces to minimize interference. However, it remains unclear how ordinal position is geometrically encoded within this framework. The position of each subspace in the ambient neural space might carry this information, as suggested by (Tian et al. 2024); alternatively, subspaces may be dynamically indexed by low-frequency oscillations, consistent with abundant evidence linking theta phase to ordinal position in sequential memory tasks. Future studies combining geometric analysis of neural subspaces with oscillatory brain activity could more directly test how interactions between neural rhythms shape subspace geometry.

Several factors in our study may also account for the observed null findings. The use of broadband time series to train classifiers within the TDLM pipeline may have limited our ability to detect neural sequences that are expressed in specific frequency bands, such as theta or gamma. Frequency-specific dynamics are thought to play a critical role in organizing sequential information, and collapsing across frequencies might obscure these patterns. However, previous MEG studies have successfully identified neural replay during resting-state periods using similar decoding approaches (Liu et al. 2019; Kurth-Nelson et al. 2016), despite the replay having well-defined spectral characteristics, suggesting that frequency specificity alone may not fully account for the absence of detectable replay in our data. Another limitation for testing the neural sequences hypothesis is the potential mismatch between the classifiers’ training data and the neural format of memory representations during the retention period. In our study, classifiers were trained on neural activity recorded during stimulus presentation, which may not capture the distinct dynamics or coding schemes that characterize mnemonic representations during delay. This misalignment could have reduced the sensitivity of the decoded state space to detect memory-related neural sequences. Future studies could address this limitation by using a memory-related functional localizer—specifically, by training classifiers on neural activity recorded during the delay period of single-item delayed match-to-sample tasks involving the same stimuli used in the main sequential working memory task. This approach may produce classifiers that are better tuned to the neural code used during memory maintenance.

In conclusion, we tested two hypotheses about the neural mechanisms underlying sequential working memory: the neural subspaces hypothesis and the neural sequences hypothesis. Our findings revealed behaviorally relevant, low-dimensional neural subspaces that organized memory representations specifically during the maintenance period. In contrast, we found no evidence of sequential neural replay during retention, arguing against the idea that maintenance relies on item-by-item reactivation. Together, these results suggest that the brain transforms temporal sequences into geometric patterns in the representational space to support efficient, stable, and behaviorally relevant maintenance of information in working memory.

## METHODS

### Sample

We recruited 29 healthy adults from the NYUAD community to participate in this study, which included two MEG sessions and one MRI session. Five participants were excluded for not completing all MEG sessions, four were excluded due to excessive motion, and three were excluded for not adhering to the fixation task (details provided below). Ultimately, 17 participants successfully completed both MEG sessions (age: 23 ± 5 [19 40] years; 7 females; time between MEG sessions: 8 ± 5 [1 18] days). Additionally, MRI data for anatomical structure was collected from 15 out of these 17 participants (two participants could not complete the MRI session due to claustrophobia). The final sample of 17 (15) participants was used for MEG analysis at the sensor (source) level. All subjects provided informed written consent in accordance with procedures approved by NYUAD’s IRB.

### Task design

For each session, subjects performed a functional localizer and a sequential working memory task (Figure 2A). Both tasks used the same set of stimuli: eight dots with a diameter of 60 pixels (1.88°), positioned at an eccentricity of 350 pixels (11°) and evenly spaced in 45° intervals around a 360° circle. During the presentation of a location, its corresponding dot turned red, while the placeholders for the other locations remained visible throughout both the stimulus presentation and the inter-stimulus interval.

In the functional localizer, each location was presented for 0.4 seconds, with an interstimulus interval of 0.2 seconds. A total of 560 stimuli were displayed in pseudorandom order, with 70 presentations per location, ensuring that no location was repeated consecutively. Participants were instructed to maintain their gaze on a fixation point. To enforce this, subjects pressed a button whenever the fixation point flickered (a 10% change in luminance), occurring randomly twelve times (fixation task). Additionally, to maintain engagement, participants pressed a button whenever the color of a dot was orange instead of red, which also occurred twelve times (color task). All participants completed both tasks successfully (fixation task: 11.2 ± 1.5 [7 12] responses; color task: 11.8 ± 0.4 [11 12]). The primary goal of the functional localizer was to collect sensory representation data for training classifiers used in the decoding analysis, independently of the working memory task.

The sequential working memory task consisted of 210 trials, divided into three blocks of 70 trials each. Unique sequences were created at random, with no consecutive repeated locations. Trials of length-3 and 4 were intertwined. The first (second) session had 115 (110) trials of length-3, and 95 (100) trials of length-4. Every trial began with a 2.2-second baseline, followed by the presentation of a sequence of spatial locations (0.3 seconds per item, with an interstimulus interval of 0.1 seconds). The sequence length varied between 3 and 4 items. After a 4-second delay, participants were prompted to reproduce the sequence using a discrete circular response controller (Current Designs, Philadelphia, PA, USA). At the end of their response, participants received dichotomous qualitative feedback. After the feedback, the next trial started only when the participant indicated they were ready by pressing a button, allowing them time to blink. Participants were instructed to keep their gaze fixed on a central point throughout the entire trial. During the delay period, they performed the same fixation task described previously, with the fixation point flickering in 10 randomly selected trials across the session, and these trials were excluded for subsequent analyses. Participants who did not successfully complete the fixation task in at least 60% of the trials in both sessions were excluded from the study (fixation task: 8.6 ± 2 [6 10] correct responses in the final sample).

### Behavioral analysis

To determine whether participants’ performance on the working memory task was significantly above chance, we conducted a binomial test for each subject and sequence length, comparing the observed number of correct responses to the expected number based on the chance levels. The probability of achieving correct responses by chance was calculated separately for sequences of length-3 (1/n^3^) and length-4 (1/n^4^), where n=8 possible locations. Additionally, we performed a linear regression analysis to assess behavioral performance. In this model, the percentage of correct responses served as the response variable, while sequence length, session, and their interaction were included as predictor variables.

### MEG and MRI acquisition

MEG data was recorded continuously using a 208-channel axial gradiometer Yokogawa system (Kanazawa Institute of Technology, Kanazawa, Japan) with a sampling rate of 1,000 Hz and an online low-pass filter of 200 Hz. During the MEG session, visual stimulation was presented in a Vpixx system (PROPixx 120Hz). An image of 1920×1080 pixels was projected on a screen of 65 x 36.5 cm, and the viewing distance was 56 cm, resulting in a viewable image of 60.26° x 36.1° of visual angle of width and height. Before MEG acquisition, each subject’s head shape was digitized using a Polhemus dual source handheld FastSCAN-II. High-resolution anatomical MRI volume was collected during a separate session to constrain the transformation from sensor to source space. The anatomical scan was acquired on a Siemens Magnetom Prisma 3T MRI scanner using a 6.5 min MPRAGE-3D T1-weighted, gradient-echo acquisition sequence (TR: 2,400 ms, TE = 2.22 ms, flip angle: 8 degrees, voxel size: 0.8 mm3, 208 slices, FOV: 256 mm).

### Preprocessing

The continuous MEG data was first noise-reduced using eight magnetometer reference channels located away from the participant’s head and using the Time Shifted Principle Component Analysis (TSPCA, block width of 5,000 and 30 shifts) as implemented in MEG160 software (Yokogawa Electric Corporation and Eagle Technology Corporation, Tokyo, Japan). All subsequent preprocessing steps were performed using FieldTrip(Oostenveld et al. 2011), according to the FLUX pipeline (Ferrante et al. 2022). Localizer data was segmented into trials based on stimulus presentation. For the working memory task, data was epoched from 1 second before stimulus onset to 4 seconds after stimulus offset, comprising baseline, stimulus presentation, and the memory delay period. Automatic artifact detection was used to exclude trials with jump, muscle and blink artifacts (z-value = 25). We kept 91 ± 3 % of trials in the localizer (valid trials: 1023 ± 30 [943 1053] out of 1120 trials across sessions), and 80 ± 10% of trials in the working memory task (valid trials: 336 ± 40 [233 380] out of 420 trials across sessions). To further clean up the data, we used Independent Component Analysis (ICA) to attenuate blink, muscle and cardiac artifacts. The noise ICs, identified by visual inspection based on their topography and timecourses, were regressed out from the data (localizer: 18 ± 4 [10 25] noise ICs; WM task: 23 ± 5 [15 30]).

Structural images were skull-stripped (FSL-bet) and registered to the MNI space with linear (FSL-flirt) and non-linear (FSL-fnirt) transformations with FMRIB Software Library (Smith et al. 2004). Then, these transformation matrices were used to compute the subject-specific parcellations of the (Schaefer et al. 2018) atlas with 200 parcellations and 7 networks. We translated sensor-level data to source space using a beamforming technique (Jaiswal et al. 2020; Westner et al. 2022). The head shape and structural MRI images were aligned to the Neuromag coordinate system, facilitating accurate coregistration between anatomical and functional data. A realistic single-shell volume conduction model was computed based on the subject’s anatomy (Nolte 2003), and the source model was generated using a 1.25 cm isotropic 3D grid inside the head model. These source and head models were subsequently used to construct a forward (leadfield) model in the native anatomical space. The resulting spatial filters were used to localize sources in the time domain with the Linearly Constrained Minimum Variance method, projecting the dipole time series in the direction of maximum variance, and computing the absolute value of the reconstructed signal to deal with sign ambiguity. Finally, we pooled sources within each atlas parcel using an eigenvariate method, and removed zero-lag correlations between parcels’ timeseries in order to address spatial leakage with symmetric orthogonalization using the MEG-ROI-nets toolbox (Colclough et al. 2015).

### Geometric analysis

The geometric analysis of neural subspaces was adapted from previous work (Panichello and Buschman 2021; Xie et al. 2022), but applied here to timeseries of broadband neural activity from 200 cortical parcels defined by (Schaefer et al. 2018). We selected 200 parcels to approximately match the dimensionality of the denoised sensor data (207 sensors minus the number of components rejected during ICA-based artifact attenuation), as increasing the spatial resolution with finer-grained cortical parcellation would lead to greater linear mixing of source time series. An overview of the geometry analysis pipeline is provided in Supplementary Figure S1. First, we constructed neural activity matrices X of size M×N, where M is the number of conditions—defined by the stimulus locations at each ordinal position in the sequence (rank)—and N is the number of channels (brain parcels). Each column of X captures the responsiveness of a given channel to the sequences presented across trials, estimated using ridge-regularized linear regression (λ = 1) applied separately to each channel. The response variable was the channel activity averaged over a predefined time window (see below), baseline-corrected by subtracting the mean activity during a 1-second pre-stimulus baseline period. The design matrix included n×r binary predictor variables, where n=8 is the number of stimulus locations and r is the rank (sequence length), indicating which stimuli were presented on each trial. The resulting beta weights for the M conditions served as the entries in that channel’s column of X. Finally, each column of the X matrix was demeaned prior to geometric analysis. We computed subject-specific X matrices separately for correct and incorrect trials. For each response type, we constructed one X matrix per sequence length (three or four items) and for each time window of interest. These time windows included the stimulus presentation period (0.3 seconds) as well as the delay period, which was divided into 13 non-overlapping windows of 0.3 seconds each. Then, we reduced the dimensionality of the X matrix along the channel dimension (columns of X) using Principal Component Analysis (using the svd.m MATLAB function applied to the covariance matrix of X). We then projected X onto the subspace defined by the top three principal components, resulting in a matrix Z of size M×3, where each row corresponds to a condition and each column to a principal component (i.e., PC scores). Then, we subset the Z matrix by sequence rank, creating Zr matrices (8 locations by r components), with r = [1 2 3] for length-3 and r = [1 2 3 4] for length-4, representing the rank-specific neural subspaces. Finally, we computed the subspace plane for each rank-specific subspace by taking the two leading components of a PCA on Z_r_. To visualize the spatial distribution of neural subspaces, we plotted brain maps of the eigenvectors corresponding to the top three principal components. For each component, we computed the absolute value of its eigenvector and averaged these across the three leading components, across time windows and across subjects. Taking the absolute value addresses the inherent sign ambiguity of PCA, preventing cancellation due to arbitrary polarity. This approach highlights the relative contribution of each brain parcel to the dominant dimensions of the neural subspaces.

To quantify the alignment between rank-specific subspaces, we computed the principal angle (PA) and variance accounted for (VAF) between subspaces (Panichello and Buschman 2021; Xie et al. 2022). PA was defined as the angle between the normal vectors of the planes corresponding to each subspace, following the normal vector approach outlined in (Panichello and Buschman 2021). VAF was calculated as the proportion of variance explained by the principal components of one rank-subspace when neural data from another rank-subspace is projected onto it, providing a measure of the similarity in neural geometry across subspaces, as described in (Xie et al. 2022). We calculated the lower (upper) bound of PA (VAF), referred to as within-subspace PA/VAF, with a bootstrapping approach. We randomly half-split trials to create two X matrices, and computed the PA and VAF between the same ranks across the two splits. We repeated the process 1,000 times and averaged across iterations. To quantify the discriminability of locations for each rank, we computed two separability metrics in the neural subspaces Z_r_. First, the euclidean distance between locations nearby in the physical space as a measure of how well the subspace captures fine-grained differences between adjacent or closely located points. Second, the volume of the set of points defining all rank-specific locations (MATLAB’s convhulln function) (de Berg et al. 2008).

We used linear regression models to perform group-level statistical analyses. Separate models were fitted for each sequence length (three or four items) and across different time windows. To test whether the alignment between rank subspaces exceeded chance levels, we fit linear models with PA or VAF as the response variable and a predictor indicating whether the comparison was between different subspaces (across ranks) or within the same subspace (split-half). To further assess whether observed alignment exceeded what would be expected from structureless data, we fit a second model with the same response variable (PA or VAF) but with a predictor distinguishing empirical data from surrogate data (see below). To assess the behavioral relevance of neural geometry, we fitted several distinct models. Each model included trial accuracy (correct vs. incorrect) as the predictor variable and one of the following as the response variable: alignment measures (PA or VAF) or separability measures (distance or volume). To examine the modulation of neural geometry by cognitive load, we applied the same set of models, but compared geometric variables between trials of length 3 vs. length 4. All *p*-values were corrected for multiple comparisons using False Discovery Rate (FDR) correction. Effects were considered significant at *p* < 0.05 (two-tailed).

To address potential bias arising from the reduced number of incorrect trials—typically around 20% of the number of correct trials—we adopted complementary strategies. First, we computed the neural subspace for incorrect trials by projecting their X matrices onto the leading principal components (eigenvectors) derived from correct trials. This yielded a Z matrix (PC scores) for incorrect trials within the same low-dimensional space as correct trials. This approach ensures that comparisons between correct and incorrect trials are made within a shared representational subspace, enabling direct assessments of geometric properties such as distances and structure. It also assumes that the neural geometry of correct trials reflects a more stable or canonical representation of task-relevant information, allowing us to quantify how much the representations during incorrect trials deviate from this normative structure. Importantly, this avoids estimating an unstable or noisy subspace from the smaller and potentially more variable set of incorrect trials, which could confound interpretation of geometric differences. Second, we implemented a controlled resampling approach to address the imbalance in trial counts. For each comparison of geometric variables, we randomly subsampled correct trials to match the number of incorrect trials, then recomputed the X matrix using only those correct trials. We projected both the subsampled correct-trial and the full incorrect-trial X matrices onto the leading principal components derived from the original correct-trial data. Geometric variables were then calculated for each iteration, and results were averaged across 1000 resampling iterations to ensure robustness.

We generated two distinct null distributions (1,000 iterations), each designed to disrupt different aspects of the data’s underlying structure: the geometry of the manifold and the variance captured by the principal components (PCs). First, to generate a null distribution of subspace alignment metrics (PA and VAF), we independently shuffled the rows and columns of the X matrices. This procedure preserves the overall covariance structure of the data—reflected in an unchanged eigenvalue spectrum—while disrupting the specific stimulus arrangement in the low-dimensional manifold. Second, to assess the dimensionality of the neural manifold, we generated a null distribution of the eigenvalue spectrum by independently shuffling the rows (conditions) within each column (channel) of the X matrix. This permutation disrupts the covariance structure between conditions and channels, yielding a covariance matrix whose eigenvalues reflect a structure dominated by noise.

We conducted several supplementary analyses to assess the robustness of our findings. First, neural subspaces were estimated by retaining the leading six and eight Principal Components instead of three, explaining 82±3% and 88±2% of the variance, respectively. For subspaces of dimensionality greater than three, the normal vector approach to computing the principal angle (PA) is not applicable, as there is no single vector orthogonal to the entire subspace. Instead, we computed principal angles between subspaces using a method based on singular value decomposition (SVD). Specifically, we first obtained an orthonormal basis for each subspace using QR decomposition. We then calculated the matrix of inner products between the basis vectors of the two subspaces and computed the singular values of this matrix, corresponding to the cosines of the principal angles between the subspaces. Because each subspace is defined as a two-dimensional plane in R^n (being n>3), this method returns two principal angles per pair of subspaces. We report both the minimum and maximum of these two angles as summary measures of subspace alignment. The choice of six and eight principal components was motivated by geometric constraints: in R^6 and R^8, respectively, it becomes possible for three and four two-dimensional subspaces to be mutually orthogonal—that is, for all pairs of subspaces to have both principal angles equal to 90 degrees. This level of orthogonality is not achievable in lower-dimensional spaces due to the overlap required between subspaces. While increasing the number of principal components expands the ambient space and theoretically allows for fully orthogonal subspaces (i.e., two principal angles of 90° between each pair of planes), this comes with an important cost. Principal angles are computed between two-dimensional planes, each defined by two vectors in R^n, where n is the number of retained principal components. These vectors are obtained as the leading two eigenvectors from a second PCA applied to the PC scores corresponding to each ordinal position (i.e., the Z_rank matrix described earlier). In lower-dimensional spaces such as R^3, the first two eigenvectors typically explain a large proportion of the variance (85±1%), making the plane a faithful representation of the underlying subspace geometry. However, as the dimensionality increases, the variance captured by the first two eigenvectors decreases (e.g., 65±2% in R^6 and 58±2% in R^8), meaning that the 2D planes used to compute principal angles become progressively less representative of the full subspace structure. As a second supplementary analysis, to mitigate the impact of limited sample size on the estimation of the X matrices, we reduced the number of stimulus locations. Rather than using all 8 locations for each ordinal position, we recomputed the X matrices using only 4 locations by grouping pairs of neighboring positions around the cardinal axes. Because each location could be grouped with either its left-side or right-side neighbor, we applied both grouping schemes and averaged the resulting matrices. This procedure yielded X matrices centered on the four cardinal locations. This analysis was motivated by the limited number of observations available to estimate the X matrix for incorrect trials. For sequences of length 3 (4), the number of trials per condition was 3±2 [range: 0–9] (5±3 [range: 0–17]), with 54% (25%) of conditions having fewer than 3 trials and 10% (3%) having no trials at all. After grouping locations, the number of trials per condition increased to 5±3 [range: 1–16] (9±5 [range: 1–29]), with only 18% (5%) of conditions having fewer than 3 trials, and no conditions with zero trials. The third supplementary analysis involved repeating the entire geometric analysis pipeline using MEG sensors (N = 207) as channels, instead of cortical parcels in source space.

### Decoding analysis and sequence effects

The analysis of neural sequences of memory representations during the delay period was performed with the temporally delayed linear modelling (TDLM) (Liu, Dolan, et al. 2021), a domain-general method for finding neural sequences. TDLM has been validated for identifying neural replay in MEG recordings of humans both at rest and task (Liu et al. 2019; Liu, Mattar, et al. 2021; Schwartenbeck et al. 2023). The TDLM process involves, first, constructing the decoded state space, and then analyzing the neural sequences within that space.

To construct the decoded state space of memory representations, we trained binomial logistic regression classifiers, one for each specific stimulus location. Classifiers were trained separately for localizer and WM task, using the 200 source-based parcels (or the 207 MEG sensors) as data features. Training was performed at each time point of stimulus presentation. To improve the signal-to-noise ratio, we downsampled the time series into 50 ms bins with 5 ms steps. The training set included: (a) positive examples, when the target stimulus was displayed on the screen, and (b) negative examples, when any other stimulus was displayed. To balance the sample sizes, negative examples were created by randomly selecting stimuli from all other locations, ensuring an equal number of negative examples for each location. The logistic regression models were L1- and L2-regularized, with lambda values of [1, 0.1, 0.01, 0.001], and classifiers accuracy was evaluated using a 5-fold cross-validation approach with 10 repetitions. For group-level statistical analysis, we calculated the area under the curve (AUC) values for each subject at every time point. To determine if the AUC values were significantly different from chance (AUC = 0.5), we performed a cluster-based permutation test. This non-parametric method was chosen for its ability to handle multiple comparisons over time points while controlling for false positives (Maris and Oostenveld 2007; Sassenhagen and Draschkow 2019). We applied a correction for multiple comparisons using a cluster-based approach, with 5000 permutations, ‘maxsum’ cluster statistic, critical z-value for identifying significant clusters was set at 2.58, and the significance threshold (alpha) was set to 0.01 (two-sided). At group level, we selected the timepoint and the regularization (L1 or L2, and lambda parameter) with the highest performance (AUC). Then, we used the subject-specific weights of the winning classifiers to predict memory reactivations during delay period, getting the decoded state space. Since the sequences presented never included all 8 locations, we defined the state space with the locations presented in each trial, resulting in a decoded state space of dimensions time by trial length. Delay period timeseries were also downsampled, averaging across bins of 50 msec in steps of 5 msec, to increase the signal-to-noise ratio. Decoding analysis was performed with MVPA-Light (Treder 2020).

The temporal structure of working memory representations was assessed on the decoded state space in a two-step linear modelling approach, as described in (Liu, Dolan, et al. 2021). In the first-level sequence analysis, we quantified the state-to-state transitions across time lags, defined as the time lapse between neural reactivations in the sequence. For each time lag delta_t, ranging from 1 to 500 ms in steps of 5 ms, we fitted the regression model

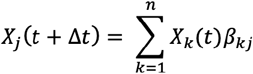

where the time-lagged copy of one state X_j(t + delta_t) was regressed onto the state space X_k(t) of the other states. When the process is completed for all states, we get a matrix of regression coefficients beta (states by states), named empirical transition matrix, defining the state-to-state transitions for the time lag delta_t. In the second-level sequence analysis, we tested the strength of particular sequences, defined by the transition matrices T_r, with a linear regression model that relates them to beta. Specifically,

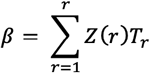

where T_r = [T_F, T_B, T_auto, T_const] was a set of predefined template matrices, including the hypothesized sequence of events running forward T_F and backward T_B, and another two matrices controlling for state autocorrelation T_auto and the average of all transitions T_const. The regression coefficients Z(r) reflects the strength of the sequences, or sequence effect. Statistical significance of forward and backward effects was tested with a non-parametric permutation-based method, creating null distributions by reshuffling the order of the states and recalculating the second-level analysis. Multiple comparisons over time lags were addressed with extreme-value or maxT correction, where the null distribution was formed by taking the maximum of sequenceness across all computed time lags for each permutation. The significance threshold across time lags was computed as the 2.5 and 97.5 percentile of the null distribution. See (Liu, Dolan, et al. 2021) for a detailed description of TDLM steps.

We tested sequence effects during the delay period using decoded state spaces generated by classifiers from both the localizer (localizer-to-delay) and the stimulus period of the working memory task (stim-to-delay). To further validate the accuracy of the analysis pipeline, we also applied the TDLM to the stimulus presentation with classifiers trained on WM stimuli period (stim-to-stim) and on localizer data (localizer-to-stim). Sequence effects were computed at trial level, then averaged across trials and subsequently across subjects. To control for finite sample effects during the first-level sequence analysis, we alternatively concatenated the trial-specific decoded state space and performed the analysis at the subject level. This approach increased the number of observations in the initial regression, aligning more closely with seminal studies on neural replay during rest (Liu et al. 2019; Kurth-Nelson et al. 2016). To account for inter-trial effects, we used linear mixed-effects model in the first-level sequence analysis to include random intercepts of trial and session. To prevent cross-trial contamination in the time-lagged decoded state, we set to zero the timepoints (rows of X) where trial boundaries were crossed. As a further step, we also tested eliminating these timepoints (rows of X) entirely, which produced equivalent results.

