## Supplementary Material for "Neural Subspaces Encode Sequential Working Memory, but Neural Sequences Do Not"

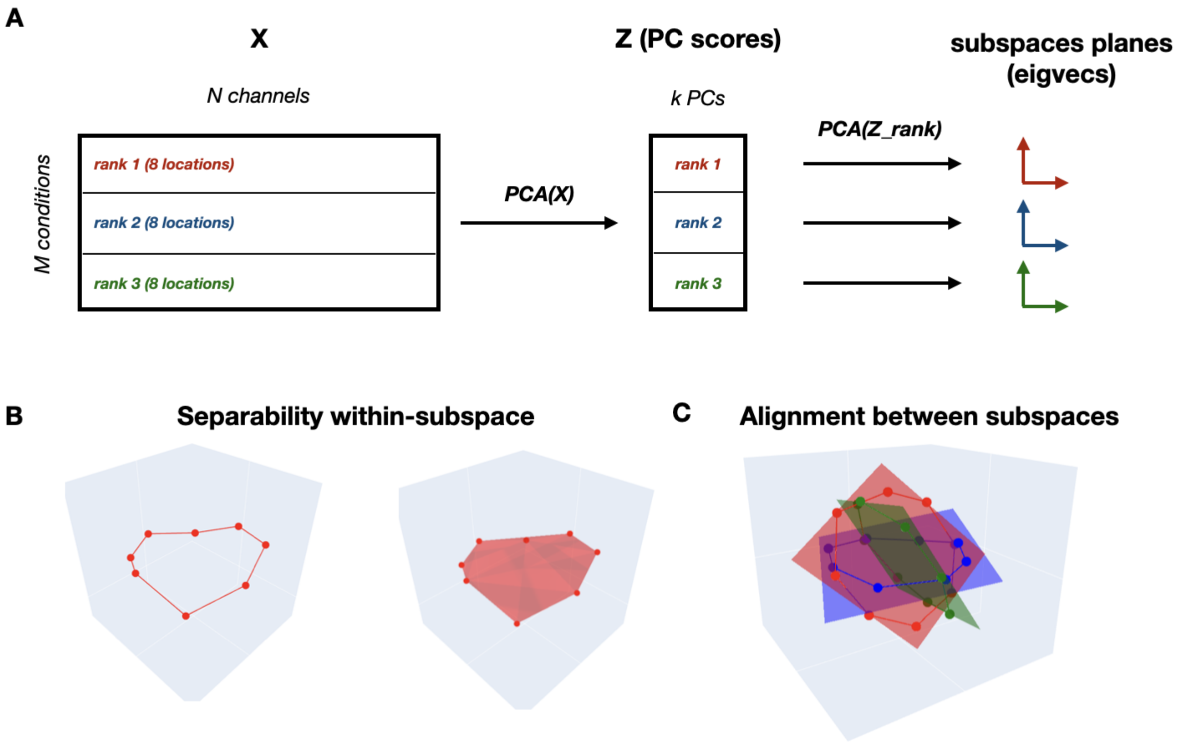


Figure S1. Geometric analysis of neural subspaces. A) Overview of the analysis pipeline. Matrix X of neural activity describes the MEG channel responsiveness to stimuli during stimuli and delay periods. Low-dimensional subspace encoding memory representations were computed with Principal Component Analysis (PCA). Projection of X matrix onto the leading k=3 PCs (PC scores) defines the neural subspace for each ordinal position in the sequence (rank-specific). A second PCA on rank-specific subspaces was applied to define the subspace planes as the two leading eigenvectors. B) Separability between stimuli representations within each subspace was computed as the euclidean distance between subspace scores (left) and the volume of the convex hull (right). C) Alignment between subspaces was computed as the Principal Angle (PA) between subspace planes, and the Variance Accounted for (VAF) between subspace scores. In B and C, we show results from a simulation in which stimuli are arranged in a noisy ring structure with 8 equally spaced locations; in C, three rings are simulated with 60-degree angular separation.

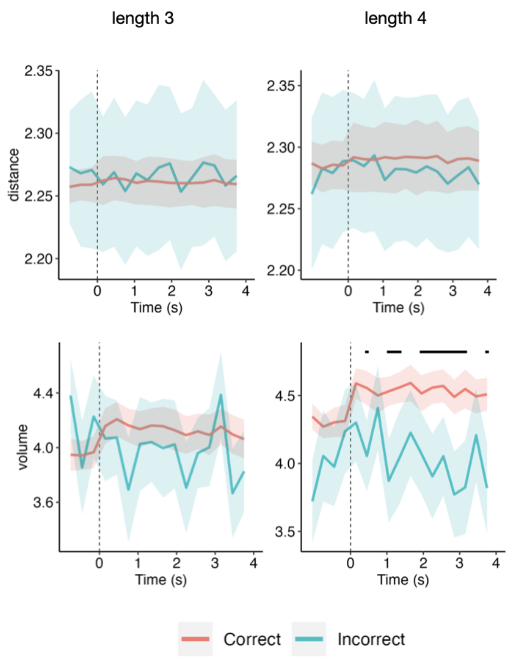


Figure S2. Geometric analysis of neural subspaces. Effect of performance on the separability of stimulus location representations in the neural subspaces, as measured by euclidean distance (top row) and volume (bottom row), for trials of length 3 (left) and length 4 (right). X-axis shows time locked at delay onset (vertical dashed line). Black horizontal line on top indicates time segments with FDR-corrected significant differences between correct and incorrect trials. Shaded areas indicate the standard error of the mean across subjects.

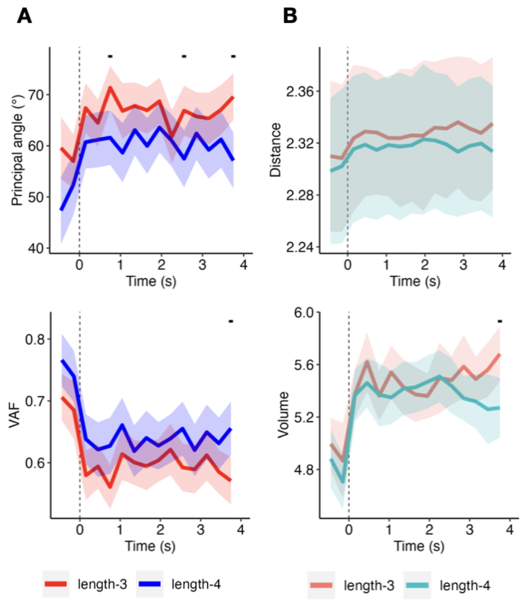


Figure S3. Geometric analysis of neural subspaces. Effect of sequence length on A) the alignment between subspaces and B) the separability of stimulus location representations in the neural subspaces. X-axis shows time locked at delay onset (vertical dashed line). Black horizontal line on top indicate time segments with FDR-corrected significant differences between correct and incorrect trials. Shaded areas indicate the standard error of the mean across subjects.

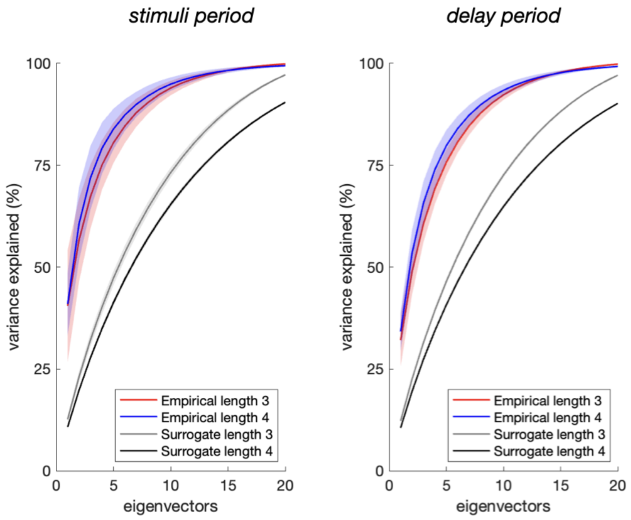

Figure S4**.** Variance explained by principal components during the stimulus period (left) and delay period (right), averaged across subjects and time windows, for trials of length 3 (red) and length 4 (blue). Gray (black) lines represent the variance explained for length 3 (4) under a null distribution of eigenvalue spectrum obtained by independently shuffling rows (conditions) within each column (channels) of the X matrix of neural activity. Shaded areas indicate the standard error of the mean across subjects.

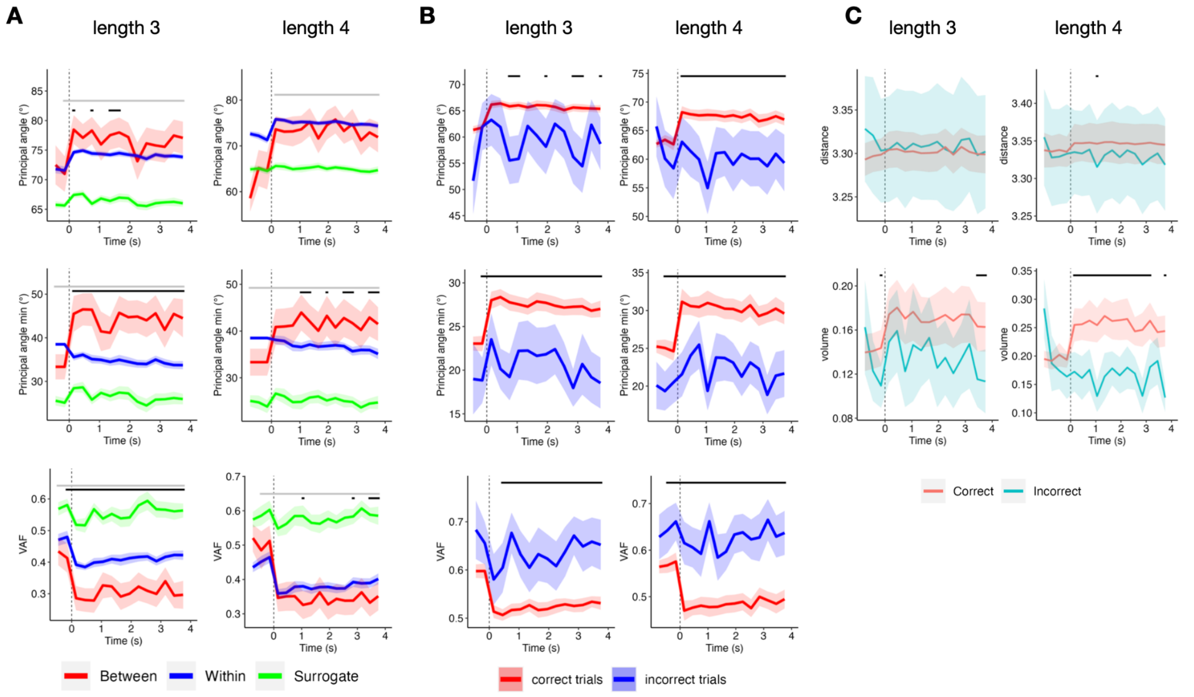


Figure S5. Geometric analysis of neural subspaces defined using the leading six Principal Components. A–B) Alignment between rank-specific subspaces, quantified by the two principal angles (PA; top and middle rows) and variance accounted for (VAF; bottom row). A) Comparison of alignment metrics for correct trials: empirical between-subspace alignment (red), surrogate between-subspace alignment (green), and within-subspace alignment (blue). B) Empirical between-subspace alignment for correct (red) vs. incorrect (blue) trials. C) Effect of performance on the separability of stimulus location representations in neural subspaces, measured by euclidean distance (top row) and subspace volume (bottom row). The x-axis shows time aligned to delay onset (vertical dashed line). Black (gray) horizontal bars indicate time segments with FDR-corrected significant differences between the variables represented by red and blue (green), respectively. Shaded areas represent the standard error of the mean across subjects.

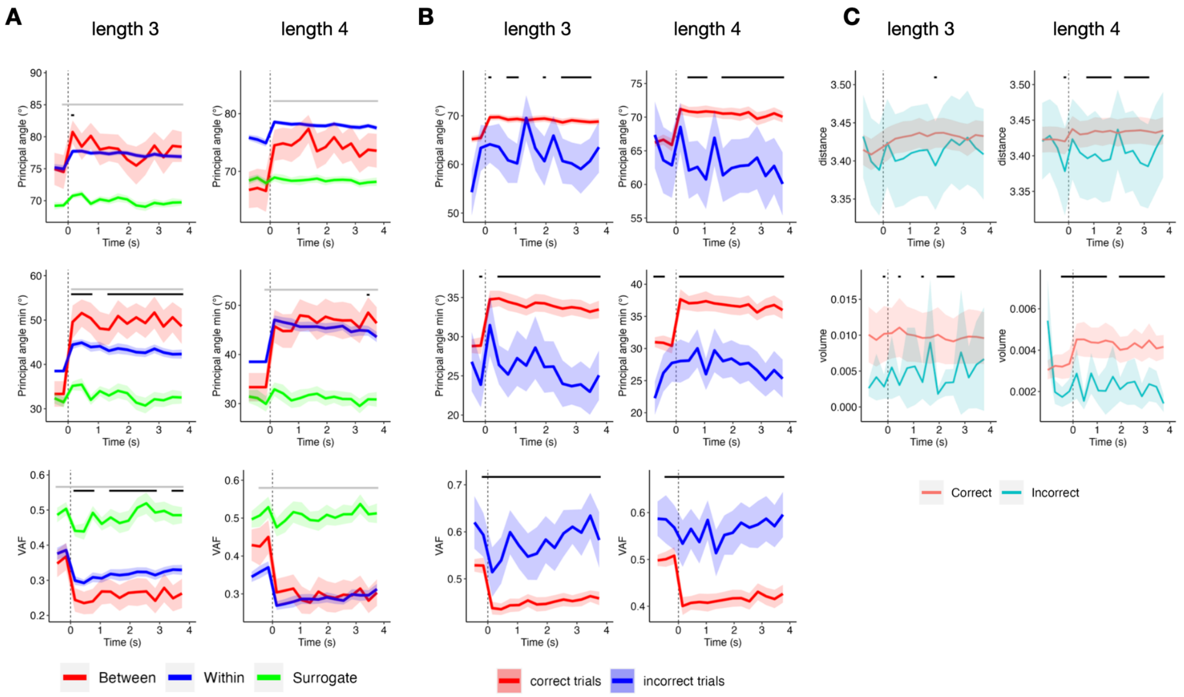


Figure S6. Geometric analysis of neural subspaces defined using the leading eight Principal Components. A–B) Alignment between rank-specific subspaces, quantified by the two principal angles (PA; top and middle rows) and variance accounted for (VAF; bottom row). A) Comparison of alignment metrics for correct trials: empirical between-subspace alignment (red), surrogate between-subspace alignment (green), and within-subspace alignment (blue). B) Empirical between-subspace alignment for correct (red) vs. incorrect (blue) trials. C) Effect of performance on the separability of stimulus location representations in neural subspaces, measured by euclidean distance (top row) and subspace volume (bottom row). The x-axis shows time aligned to delay onset (vertical dashed line). Black (gray) horizontal bars indicate time segments with FDR-corrected significant differences between the variables represented by red and blue (green), respectively. Shaded areas represent the standard error of the mean across subjects.

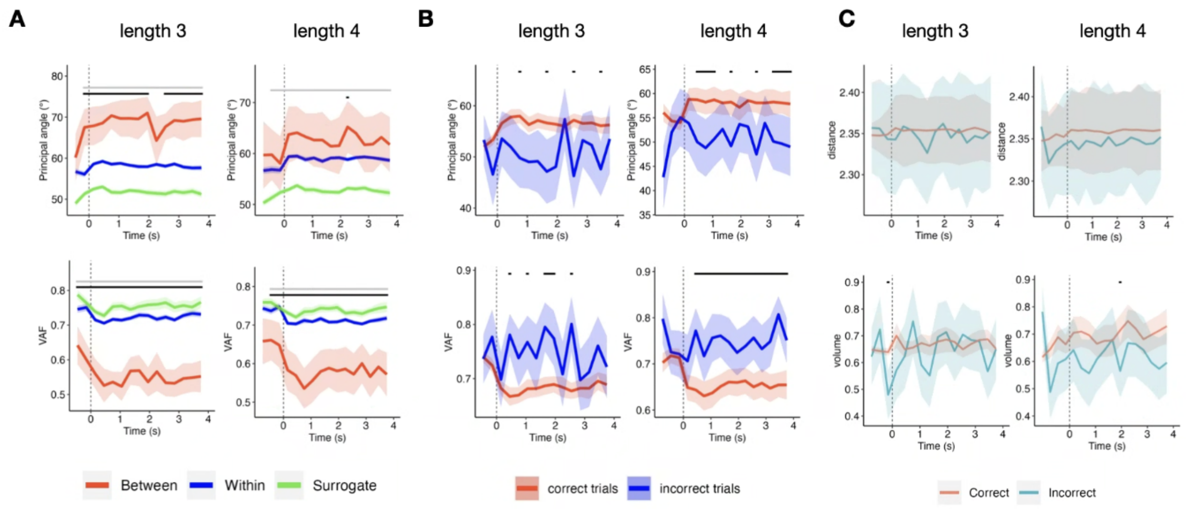


Figure S7. Geometric analysis of neural subspaces when the number of spatial locations in the matrices of neural activity were downsampled to increase the signal-to-noise ratio. A–B) Alignment between rank-specific subspaces, quantified by the two principal angles (PA; top rows) and variance accounted for (VAF; bottom row). A) Comparison of alignment metrics for correct trials: empirical between-subspace alignment (red), surrogate between-subspace alignment (green), and within-subspace alignment (blue). B) Empirical between-subspace alignment for correct (red) vs. incorrect (blue) trials. C) Effect of performance on the separability of stimulus location representations in neural subspaces, measured by euclidean distance (top row) and subspace volume (bottom row). The x-axis shows time aligned to delay onset (vertical dashed line). Black (gray) horizontal bars indicate time segments with FDR-corrected significant differences between the variables represented by red and blue (green), respectively. Shaded areas represent the standard error of the mean across subjects.

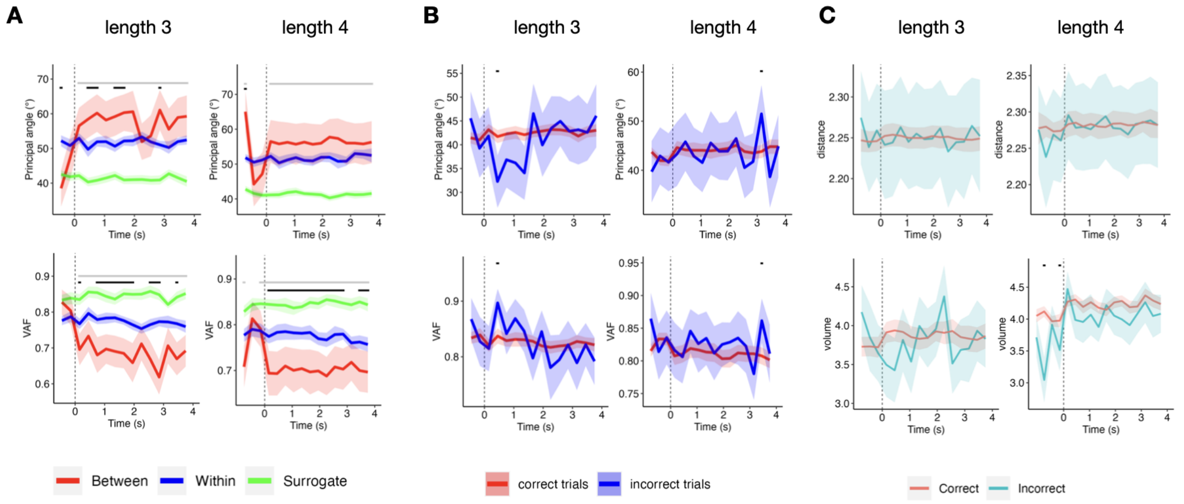


Figure S8. Geometric analysis of neural subspaces when carried out in the sensor space. All figure specifications and analysis procedures are identical to those described in Figure S7.

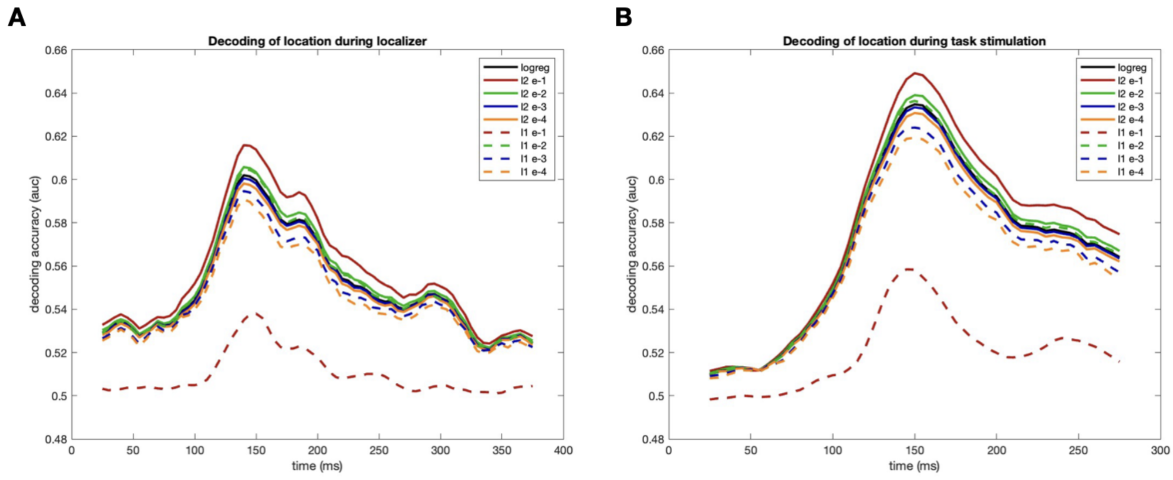


Figure S9. Decoding performance of stimulus location on stimulus presentation during localizer (A) and sequential WM task (B) with all classifiers. Lines show the average of area under the curve (auc) across subjects. Logreg, logistic regression, l1, L1 regularization; l2, L2 regularization; e-n, lambda value of regularization, with n = [1 2 3 4]. All classifiers were significant around the peak at 150 ms, except L1e-1.

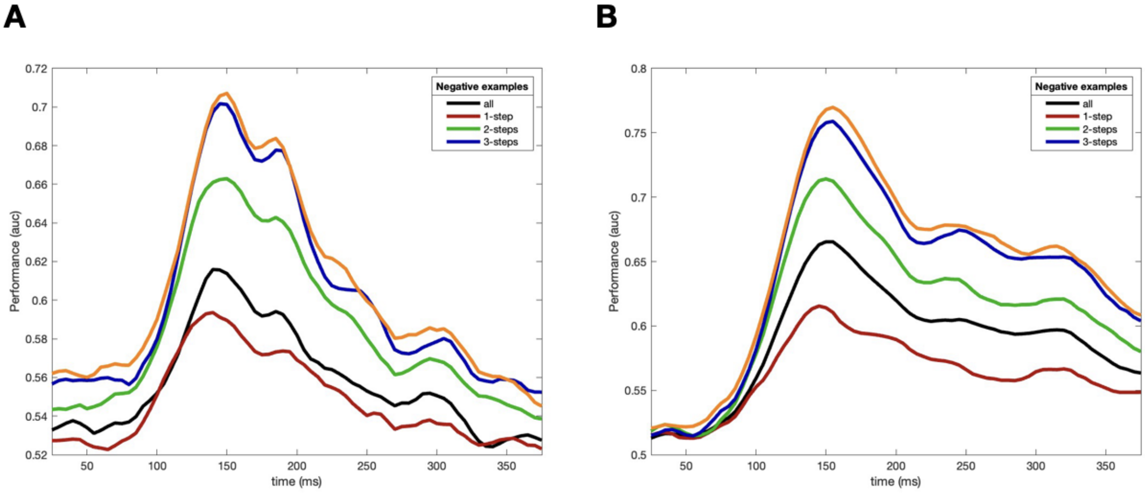


Figure S10. Decoding accuracy of source-space L2 e-1 classifiers (AUC, y-axis) as a function of time after stimulus onset (x-axis) during (A) localizer and (B) WM task. Different lines indicate the subset of locations used in the negative examples of the training set. Black line indicates classifiers performance when all stimuli were included in the training set. Colored lines indicate performance when negative examples in the training set only included locations at a distance of n-steps to the positive example. For example, when the training set contained location 4 as positive example, 1-step indicated that negative examples included locations 3 and 5; while 2-steps indicated that negative examples included locations 2 and 6.

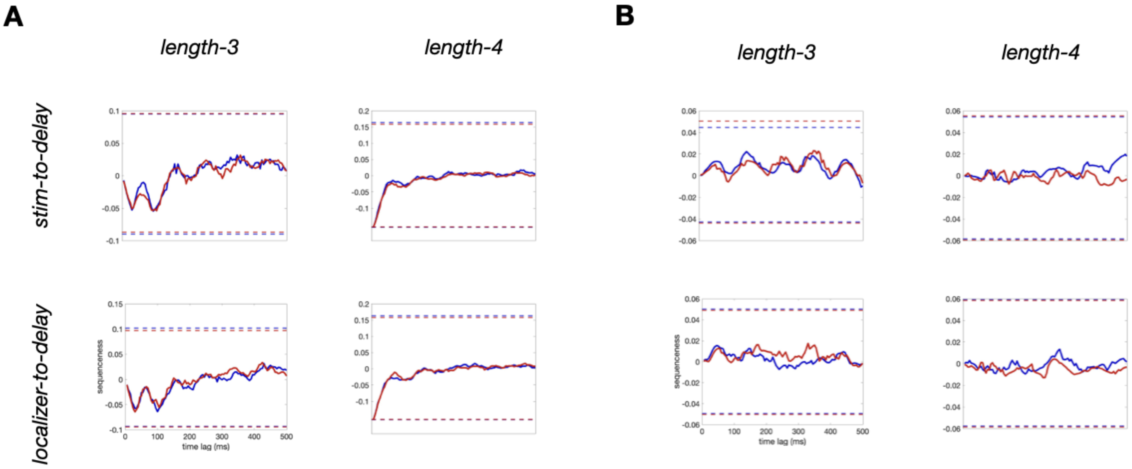


Figure S11. Sequenceness effects computed with the subset of trials without locations at 1-step distance, computed at (A) trial-level and (B) subject-level. Rows (top to bottom) show: a) stim-to-delay: classifier trained on WM-stimuli data and applied on WM-delay data, and b) localizer-to-delay: classifier trained on localizer data and applied on WM-delay data. Solid lines represent forward (blue) or backward (red) sequenceness effects averaged across subjects, dashed lines indicate the upper and lower thresholds for statistical significance.
